# Spatial Expression Pattern and Cellular Organization of Gap Junctions in Third Instar Wing Imaginal Discs of *Drosophila melanogaster*

**DOI:** 10.1101/2025.09.30.673880

**Authors:** Spraha Bhandari, Ankita Chodankar, Franka Eckardt, Reinhard Bauer

**Affiliations:** National Centre for Biological Sciences, Tata Institute of Fundamental Research, Bangalore 560065, India; Department of Biology, University of Florida, United States of America, U.S.A; Molecular Developmental Biology, LIMES-Institute, University of Bonn, Germany

**Author notes:** Corresponding author (s):, (RB).

## Abstract

The *Drosophila* wing imaginal disc serves as a powerful model to study intercellular communication during development. In our study, we report and discuss the expression pattern and cellular distribution of innexin-1, innexin-2 and innexin-3 in the cells of the wing imaginal discs. Our immunohistochemical data show that all three innexins are broadly expressed across the membranes of both the disc proper and peripodial epithelial cells of the wing disc. The stainings further reveal that, within the disc proper epithelium, junctional proteins are arranged in a clear apico-basal hierarchy: cadherins at the apical surface, followed by septate junction proteins, with innexins localised sub-apically beneath these components. All three innexins are enriched within this sub-apical domain, and are additionally detected at mid-and baso-lateral sites in varying levels. Notably, innexin-2 exhibits partial colocalization with coracle, a septate junction–associated protein, suggesting a functional association. In the peripodial epithelium, innexins are detected in distinct punctate patterns across cell membranes, implying heterogeneity in their molecular characteristics. To validate these expression patterns, we carried out tissue-specific RNAi-mediated knockdowns using the *pannier-Gal4* driver targeting the notum, a structurally and functionally important but underexplored region in innexin research. Knockdown of *innexin-2* and *innexin-3* led to complete loss of their expression within this region. Notably, silencing of *innexin-2* also affected the expression of septate junction associated proteins and *innexin-3* knockdown was accompanied by a significant reduction in disc size and altered morphology. These findings depict and confirm the presence of innexins in the notum region and also indicate that individual innexins may have distinct or shared functional roles within the same tissue domain of expression. Their localization to specific membrane domains is likely to underlie their differential modes of action. Although previous studies have demonstrated the functional involvement of gap junctions in various aspects of normal wing development in *Drosophila*, a description of the arrangement of innexins on the third instar wing discs is required for better understanding of their roles. Our study addresses this gap by providing a comprehensive analysis of the cellular localisation and organisation of gap junctions, specifically innexin-1,-2 and-3, within the third instar wing discs, thereby supporting and extending existing knowledge.

## Introduction

In *Drosophila*, the wing imaginal discs are sac-like epithelial precursor organs that originate from a cluster of 40-50 epithelial cells in the embryo (Bate & Arias., 1991). Through proliferation, they develop to form a structure of 40,000-50,000 cells by the larval stages (Madhavan & Schneiderman, 1971; García-Bellido & Merriam, 1971). During metamorphosis, the cells from the wing discs differentiate, contributing to the formation and patterning of the adult cuticle, wings, hinge, thorax epithelium and muscles of a fly (Waddington H Conrad, 1940; Fristrom & Fristrom, 1993; Martin-Blanco E *et al*., 2000; Usui & Simpson, 2000; Kanca *et al*., 2014; Kiger JA Jr *et al*., 2007). Structurally, the wing disc epithelia comprise of two types of cells, namely, the flat squamous cells forming the peripodial epithelium and the columnar cells of the disc proper. In addition, the wing discs also function as a niche for the adult muscle progenitors or adepithelial cells, trachea, neuron and glia cells etc. (Gunage *et al.,* 2014; Inoue & Hayashi, 2007; Hayashi & Kondo, 2018; Giangrande, 1994; Huang *et al*., 1991). From a scientific perspective, wing discs are valuable model systems for understanding several biological processes like pattern formation, signalling, growth, regeneration, intercellular communication in intricate tissues, injury, cancer development, wound healing etc. Gap junctions in invertebrates are formed by a family of proteins known as innexins. Through the formation of intercellular channels, innexins, like their vertebrate counterparts, called connexins, enable direct electrical and metabolic coupling critical for tissue architecture and coordination. Among the eight *innexins* identified in *Drosophila*, *innexin-1* is essential for normal optic lobe development and plays a critical role in the post-embryonic development of the central nervous system (Lipshitz & Kankel, 1985; Holcroft *et al*., 2013). Innexin-2, the most frequently observed member of this family, is the smallest (42.58 kDa) and can assemble into heteromeric channels with Innexin-1 and Innexin-3 across various tissues (Lehmann *et al*., 2006; Richard & Hoch, 2015; Richard *et al*., 2017). Innexin-2 is implicated in multiple biological processes, including morphogenesis - embryonic epithelia (Bauer *et al*., 2004), foregut (Bauer *et al*., 2002), and CNS development (Ostrowski *et al.,* 2008); organ size regulation (Holcroft *et al*., 2013; Richard & Hoch., 2015; Richard *et al.,* 2017); transfer of metabolites to regulate growth and development (Ayukawa *et al*., 2012; Boumard *et al*., 2025; Vachias *et al*., 2025; Starich *et al*., 2020) etc. Previous studies also indicate that innexins-1,-2 and –3 exhibit overlapping expression in embryonic epithelium, with innexin-3 functioning either as hemi-channels or in heteromeric association with innexin-2 (Stebbings *et al*., 2000). Furthermore, innexin-3 is essential for proper dorsal closure during embryogenesis (Giuliani *et al*., 2013). Interestingly, mRNA *in-situ* hybridization studies have demonstrated the expression of *innexin* genes within the wing discs of *Drosophila* (Stebbings *et al*., 2002). In the past, reports regarding the presence of gap junctions in the wing discs were described utilising the dye injection (Weir & Lo, 1982), mRNA *in-situ* hybridisation (Stebbings *et al*., 2002) and scanning electron microscope (SEM) based methods (Ryerse & Nagel, 1984). Additionally, experiments on the adult wing tissues have identified gap junctions through polymerase chain reaction (PCR) and RNA-seq based approaches (Raad & Robinchon, 2019; Agnel *et al*., 2017). More recently, innexin-2 was shown to exhibit a distribution pattern consistent with its mRNA *in-situ* expression, being present throughout the discs and facilitating nucleotide pool sharing between adjacent cells (Boumard *et al*., 2025). Functionally, gap junctions in the wing disc epithelia and adult wing tissue have been reported to play crucial cellular and biological roles like metabolite transport (Ayukawa *et al*., 2012; Boumard *et al*., 2025), injury response (Restrepo & Basler, 2016), regulation of growth (Jursnich VA *et al*., 1990) and organ size (Soundarrajan *et al*., 2021) and maturation of functional wing chemosensilla (Raad & Robinchon, 2019). These experiments establish both the presence and functional significance of gap junctions in multiple aspects of *Drosophila* wing development and highlight the need to further define the spatial distribution of innexins in third instar wing imaginal discs. Based on the current knowledge, this study is the first to detail the cellular localization and organisation of gap junctions in the wing imaginal discs of the third instars, thereby corroborating and expanding previous findings. In this manuscript, we specifically report the expression pattern and cellular distribution of innexin-1,-2 and-3 in the cells of the wing disc epithelia. Through our experiments, we observe that all three innexins are present throughout the disc and are predominantly localised to the membranes of the epithelial cells, where they exhibit a continuous and punctate distribution pattern. Our investigations further show that, innexins are positioned beneath the adherence and septate junction proteins along the apico-basal axis of the disc proper cells of the wing disc. Although, all three innexins are enriched at the apico-lateral domains (placed sub apically), innexins are additionally detected at mid-and baso-lateral sites. This is in contrast to eye discs, where innexins are restricted to the apico-lateral domains, underscoring tissue-specific differences in their spatial organisation. To further validate the expression patterns, we performed RNAi-mediated knockdown of *innexin-2* and *innexin-3* using a tissue-specific Gal4 driver targeting the notum region of the wing disc.

Notably, *innexin-3* knockdown led to its complete loss in the notum and was associated with a significant change in shape of cells and an overall reduction in disc size, indicative of its role in regulating epithelial cell polarity and tissue size. Their down regulation also disrupted the membranes of epithelial cells on the basal side, significantly altering the morphology of the discs. Given the localization of innexin-3 towards the basal side as well, its knockdown may further compromise basement membrane integrity; however, this possibility requires additional investigation. In contrast, *innexin-2* knockdown did not visibly affect disc size much; however, besides a reduced innexin-2 expression in the targeted region, the expression of coracle, a septate junction–associated protein is diminished. These observations indicate that *innexin-2* is critical for maintaining epithelial tissue integrity and likely operates in coordination with other membrane and membrane-associated proteins. Collectively, our immunostaining analyses define the spatial cellular distribution of innexins in the wing disc, and functional knockdown of co-expressed *innexin-2 and-3* demonstrate and indicate their likely involvement in regulating cellular and developmental processes during wing disc morphogenesis.

### Experimental Procedures

1. Drosophila culturing and fly stocks utilised in the study: All *Drosophila melanogaster* strains were reared on standard cornmeal-based medium at 25°C, unless specified otherwise. The wild-type strain Canton-S was used as a control for all experiments. For genetic knockdown experiments, the UAS-RNAi lines targeting the *innexin-2 (UASwizInx2)* and *innexin-3 (UAS-RNAi-Inx3)* were employed (Lechner *et al.,* 2007; Lehmann *et al.,* 2006). The *pannier-Gal4* driver line, *pnr-Gal4/TM6B,Dfd-EYFP,Sb,* used for tissue-specific knockdown of *innexins* was derived from the parent line procured from the Bloomington Drosophila Stock Center (BDSC: 3039). Additionally, the recombinant fly lines, *pnr-Gal4,UAS-mCD8::GFP*/*TM6B,Tb* and *pnr-Gal4,UAS-mCD8::GFP*/*MKRS*, were generated in-house to validate Gal4 activity and visualize the pannier expression domain.
2. Generation of recombinant fly line: Recombinant *pnr-Gal4, UAS-mCD8::GFP* lines were generated via meiotic recombination between *pnr-Gal4/TM6B,Tb* and *Pin/CyO; UAS-mCD8::GFP* stocks. Non-Tb F1 progeny were crossed to *If/CyO; MKRS/Tb* balancer virgins, and F2 progeny carrying both *pnr-Gal4* and *UAS-mCD8::GFP* were identified by GFP fluorescence. Stable stocks were maintained on balancers and verified across two generations using GFP expression and immunohistochemistry.
3. RNAi experiments: Tissue-specific gene knockdown was achieved by crossing males from the *UAS-RNAi* lines targeting *innexin-2* and *innexin-3* with virgin females carrying the *pnr-Gal4/TM6B,Dfd-EYFP,Sb* driver, enabling RNAi expression specifically in the notum region of the wing imaginal discs. All crosses were maintained at 25 °C and, non-EYFP larvae were selected for wing disc dissections.
4. Immunohistochemistry of the wing discs: For immunohistochemistry experiments, the wing discs were dissected from the wandering third instar larvae (120hr onwards) and then fixed in 4% paraformaldehyde solution (prepared in 1X PBS, pH 7.5) for 10-15 minutes. Following this, the tissue was subjected to three washes of 0.1% PTX (1X PBS + 0.1% Triton-X) for 10 minutes each. Subsequently, the wing discs were incubated with 10% goat serum (prepared in 0.1% PTX) for 2hrs at room temperature. Primary antibody staining was then performed by incubating the dissected discs with the appropriate antibodies for overnight at 4°C on a shaker, followed by three washes with 0.1% PTX (1X PBS + 0.1% Triton-X) for 10 minutes each. After the washes, the tissue was incubated with secondary antibody mixture for 3 to 4 hrs on a shaker at room temperature. The excess of unbound secondary antibodies were removed by three washes of 0.1% PTX (1X PBS + 0.1% Triton-X) 10 minutes each prior to mounting in Vectashield mounting media. For immunostainings, Anti-Inx-1, Anti-Inx-2 and Anti-Inx-3 (Rabbit, 1:50, kindly provided by Alexander Borst, MPI, Munich, Germany), Anti-coracle (Guinea Pig, 1:500, kindly provided by Richard Fehon, University of Chicago, USA) and Anti-dCAD2 (Rat, 1:20, DSHB) primary antibodies were used. The secondary antibodies from Invitrogen conjugated with Alexa Fluor 488, 568 and 647 were used in a ratio of 1:500 in all staining procedures.
5. Confocal data collection: Fluorescent images were captured using the Olympus FV3000 upright laser-scanning confocal microscope and processed using the ImageJ and Fiji software. All datasets were captured in multi-channel mode, with a z-step interval of 0.3 μm between optical sections. Identical laser settings were maintained across both the control and experimental samples to ensure consistency. Confocal data were acquired using 20X air and 60X oil objectives.

## Results

1. Expression pattern of innexin-1, innexin-2 and innexin-3 in the wing imaginal discs:

The third instar wing imaginal disc consists of two epithelial layers: the columnar disc proper (DP) and the overlying squamous peripodial epithelium (PE). Using the anti-innexin-1,-2 and-3 antibodies, we detected their expression throughout the entire disc, spanning both the epithelial layers (Figure 1). This expression pattern aligns with previous mRNA *in-situ* hybridization data for innexin-1 and innexin-2, but differs with innexin-3, whose transcript is not detected in the notum (Stebbings *et al*., 2002). To more precisely determine the localisation of innexin channel proteins along the apico-basal axis of the disc proper epithelium and to assess their distribution in the peripodial layer, we carried out co-immunostaining with the established membrane and cell polarity markers (Müller 2000). Since the apical surfaces of disc proper and peripodial cell layers appose each other, anti-DEcadherin antibodies were used to mark the adherens junctions on the apical side of the cells and Anti-coracle antibodies were used to label the septate junctions on the apico-lateral region of epithelial cells (Tepass and Hartenstein, 1994; Tepass *et al*., 2001). Coracle is a septate junction associated protein and a member of the Protein 4.1 superfamily of cytoplasmic proteins (Fehon *et al*., 1994).

**Figure 1.**
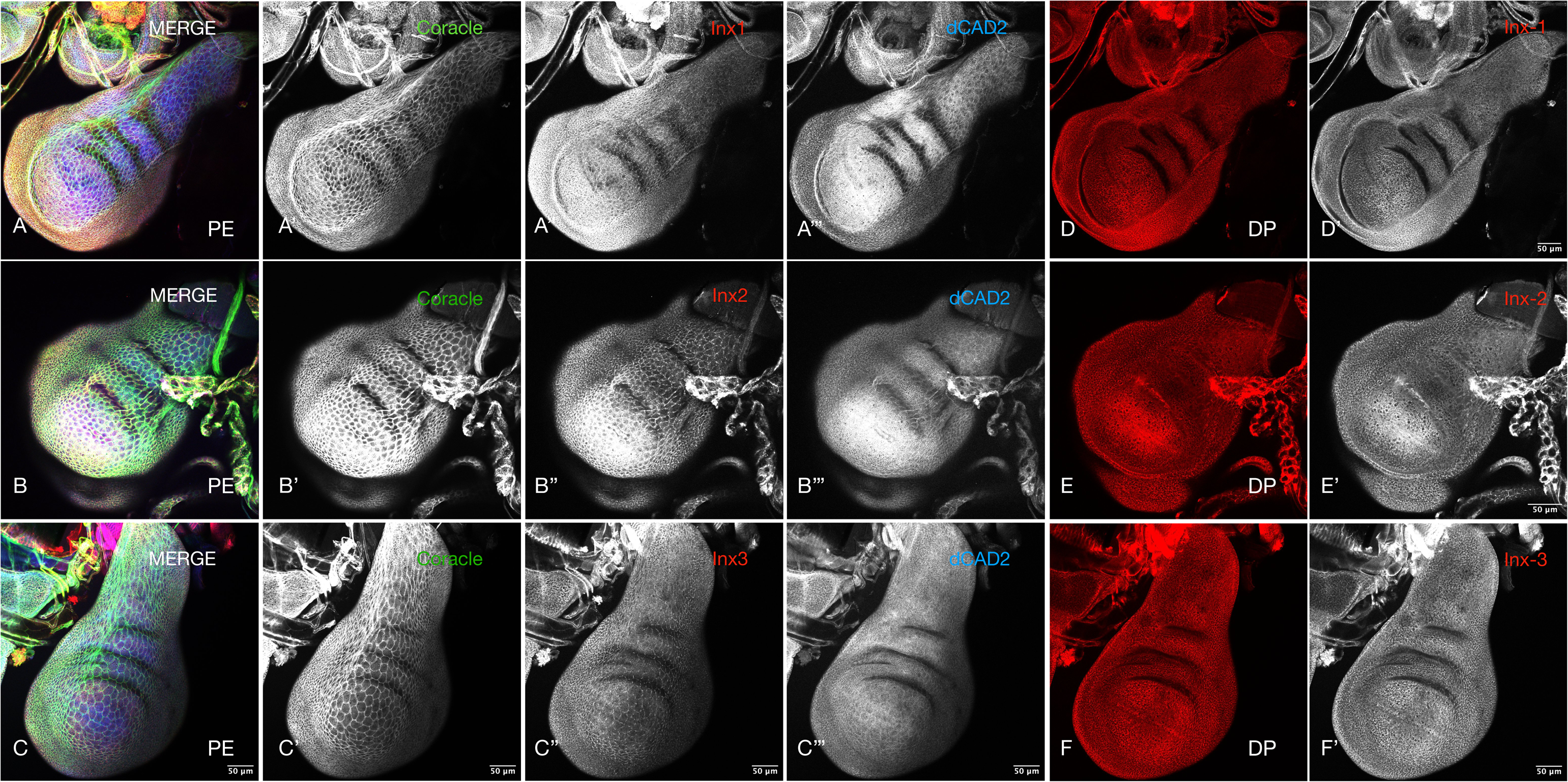
Expression pattern of innexins in the peripodial and disc proper epithelia of third instar wing imaginal discs. (A-C) Confocal images of the wing discs immunostained for Coracle (green), Innexin (red), and Cadherin (blue). **(A’-A’’’, B’-B’’’ and C-C’’’)** Corresponding grayscale images showing individual channels. **(A’’, B’’ and C’’)** Innexin-1,-2, and-3 display punctate membrane localization in peripodial cells. **(DD’, EE’ and FF’)** Color and grayscale images showing innexin expression in the disc proper epithelium. Innexins are expressed throughout the disc, including the notum, hinge, pouch, and margin regions. Wing discs were isolated from 5–6days old larvae and immunostained with antibodies against the indicated markers. **Scale bars: 50 μm. PE: Peripodial Epithelium, DP: Disc Proper**

Analysis using these markers along with innexin antibodies revealed that all three innexins were localised to the membranes of both epithelial cell types in a continuous as well as punctate pattern. Through our immunostainings, we observe that, in the peripodial epithelium, innexins exhibit distinct punctate localization patterns: innexin-1 appears as fine, scattered puncta of weak intensity, innexin-2 forms larger, more conspicuous clusters that partially overlap with septate junction markers, and innexin-3 is organized as continuous stretches of puncta along the membranes (Figure 2). Interestingly, within the peripodial epithelium, adherens and tight junctions have been described at both bicellular and tricellular interfaces (Bosveld & Bellaïche, 2020). However, at the resolution examined here, innexins were enriched at bicellular contacts, suggesting roles in intercellular communication through short-and long-range signaling or in facilitating metabolite exchange (Figure 2). Comparable punctate staining patterns have been described for innexins in other model systems (Bauer *et al*., 2004; Bohrmann & Zimmermann, 2008) and for connexins, including discrete puncta distributed across oligodendrocyte cell bodies and processes in the brain (Rash *et al., 2001*), as well as variable punctate localizations of Cx37, Cx40, and Cx43 in the murine renal vasculature (Zhang & Hill, 2005), implying that such punctate organization is a characteristic feature of gap junction proteins.

**Figure 2.**
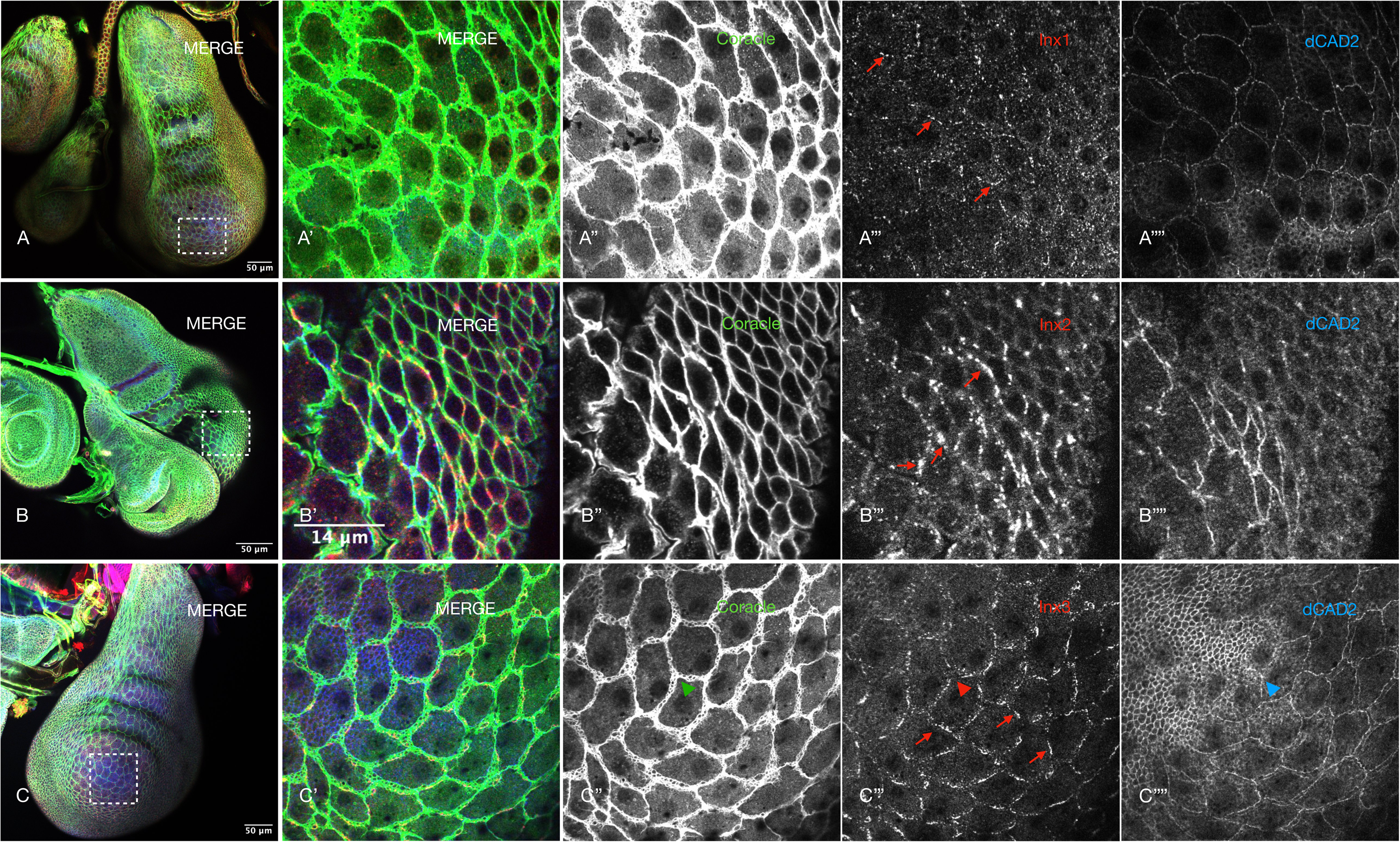
Distinct punctate localization patterns of innexins at peripodial cell membranes. (A–C) Confocal images of third instar wing imaginal discs immunostained with antibodies against Coracle (green), Innexin (red), and Cadherin (blue) proteins. **(A’–A’’’’)** Enlarged view of the boxed region in A, showing innexin-1 distributed as small, lightly stained puncta **(A’’’)** along the cell membranes. **(B’–B’’’’)** Enlarged view of the boxed region in B, depicting innexin-2 as bright, discontinuous puncta **(B’’’)** localized at the membranes. **(C’–C’’’’)** Enlarged view of the boxed region in C, highlighting Innexin-3 forming small but continuous puncta **(C’’’)** along the membranes. Red arrows indicate the specific punctate localization of each innexin subtype. Coloured triangles in C’’, C’’’ and C’’’’ indicate the tricellular junctions, where innexins are less prominent. Scale bars: 50 μm (A–C) and 14 μm in higher magnification panels (A’-A’’’’, B’-B’’’’ and C-C’’’’).

Confocal imaging of the disc proper further demonstrated that innexins localize at both the apical and basal sides of the wing disc (Figure 3). More specifically, towards the apical side, innexins localise to a sub-apical (apicolateral) domain, positioned directly below adherens and septate junction proteins. Although all three innexins shared this general sub-apical distribution, with prominent enrichment at apicolateral sites, innexins are also present at the mid and basolateral sites. Innexin-1 was relatively sparse toward the basal region, whereas innexin-2 and innexin-3 extended further into the mid-and baso-lateral sites of the disc proper epithelium (Figure 4). This arrangement is different from that observed in eye discs, where innexins are localised to the apico-lateral domains towards the apical side only. Within the disc proper, innexin-2 also exhibits partial colocalization with the septate junction-associated protein, coracle (Figure 4B), suggesting a potential functional interaction. This distribution contrasts with that observed in other polarized epithelial systems, such as the embryonic and larval body wall epithelia where the expression domain of innexin-2 is apical to that of coracle, and colocalized with Armadillo and DE-cadherin (Bauer *et al*., 2004; Luedke *et al*., 2024). Additional distinctions are evident in hindgut and salivary gland epithelia, where innexin-2 is localised to lateral domains in the hindgut and basolateral regions in salivary gland cells (Bauer *et al*., 2003; Bauer *et al*., 2005). A similar phenomenon is observed in vertebrate systems, where different connexins, functional homologs of innexins, occupy specific membrane subdomains. For example, In thyroid epithelial cells, Cx43 localizes subapically near the tight junction belt, while Cx32 forms punctate clusters along the lateral membrane (Guerrier *et al*., 1995). Under pathological conditions, connexin organization is altered, with Cx26 and Cx43 redistributed toward basolateral regions compared to their normal apicolateral and basolateral positions (Al-Ghadban et al., 2016). The broad distribution of gap junction proteins in the wing disc, distinct from patterns seen in other polarized tissues, highlights their likely contribution to morphogenetic processes by facilitating epithelial cell shape transitions crucial for wing disc development (McClure and Schubiger, 2005).

2. Genetic knockdown of *innexins* in the notum region:

**Figure 3.**
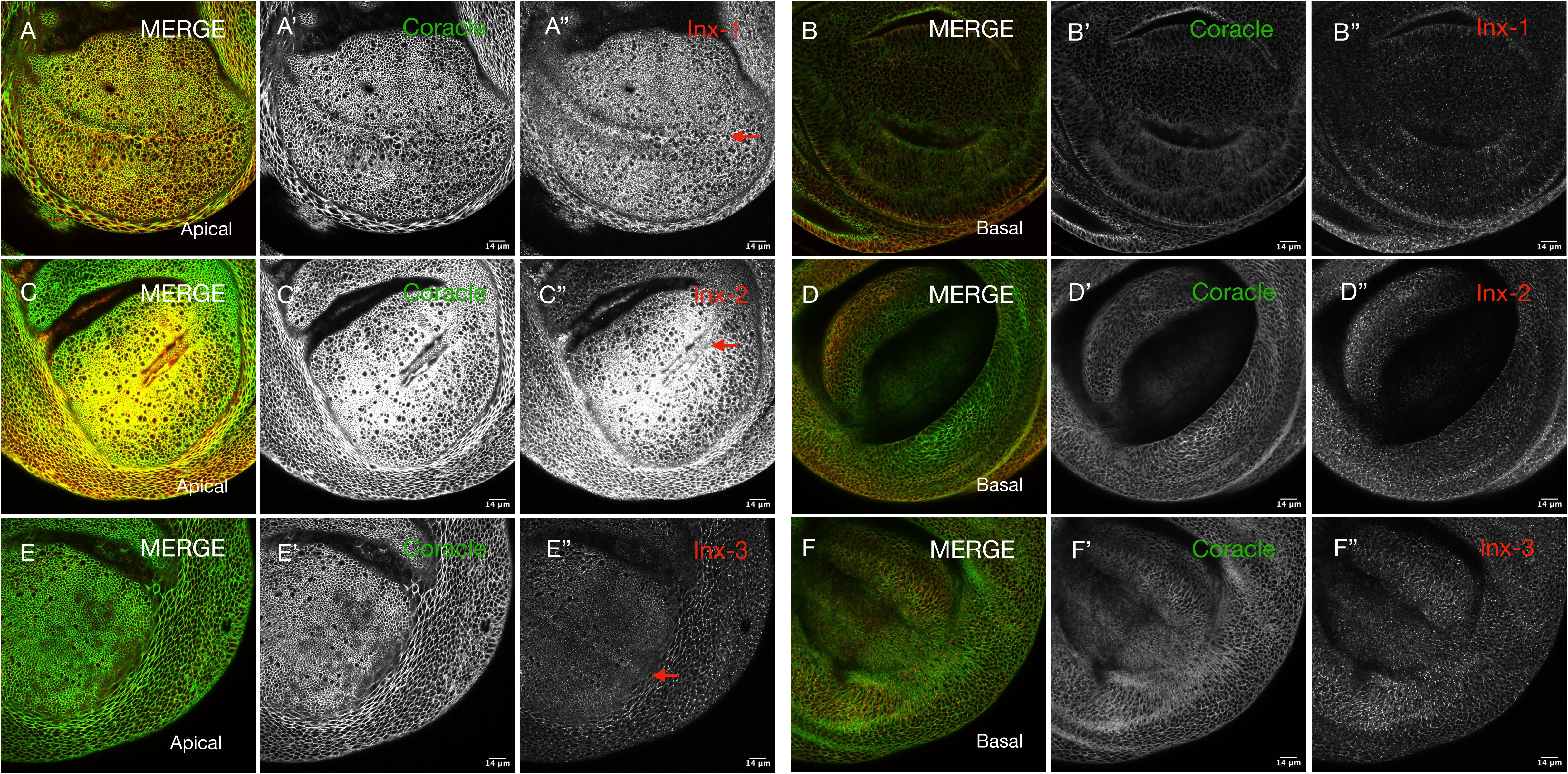
Apical and basal distribution of innexin-1,-2, and-3 in the wing disc. Confocal Z-sections of the wild-type wing disc pouch, stained with anti-innexin (red) and anti-Coracle (green) antibodies, reveal innexin localization on both the apical **(Panels A, C, E)** and basal **(Panels B, D, F)** sides of the disc proper epithelium. Unlike in the eye disc, where innexins are confined to apico-lateral domains, all three innexins in the wing disc are broadly distributed, with enrichment at the apical surface. Notably, innexin-3 also shows strong enrichment at the basal side. **Panels A, C, and E** highlight the apical distribution of innexin-1,-2, and-3, respectively. Innexin-2 displays particularly high accumulation in the pouch region **(C’’)**, and all three innexins are enriched at the central boundary of the pouch **(red arrows in A’’, C’’, D’’)**, in addition to being broadly expressed throughout the pouch. Comparative views of the apical **(A’’, C’’, D’’)** and basal **(B’’, D’’, F’’)** planes further demonstrate that innexin-3 exhibits comparable expression levels on both sides of the epithelium. **Scale bars: 14 μm**

**Figure 4.**
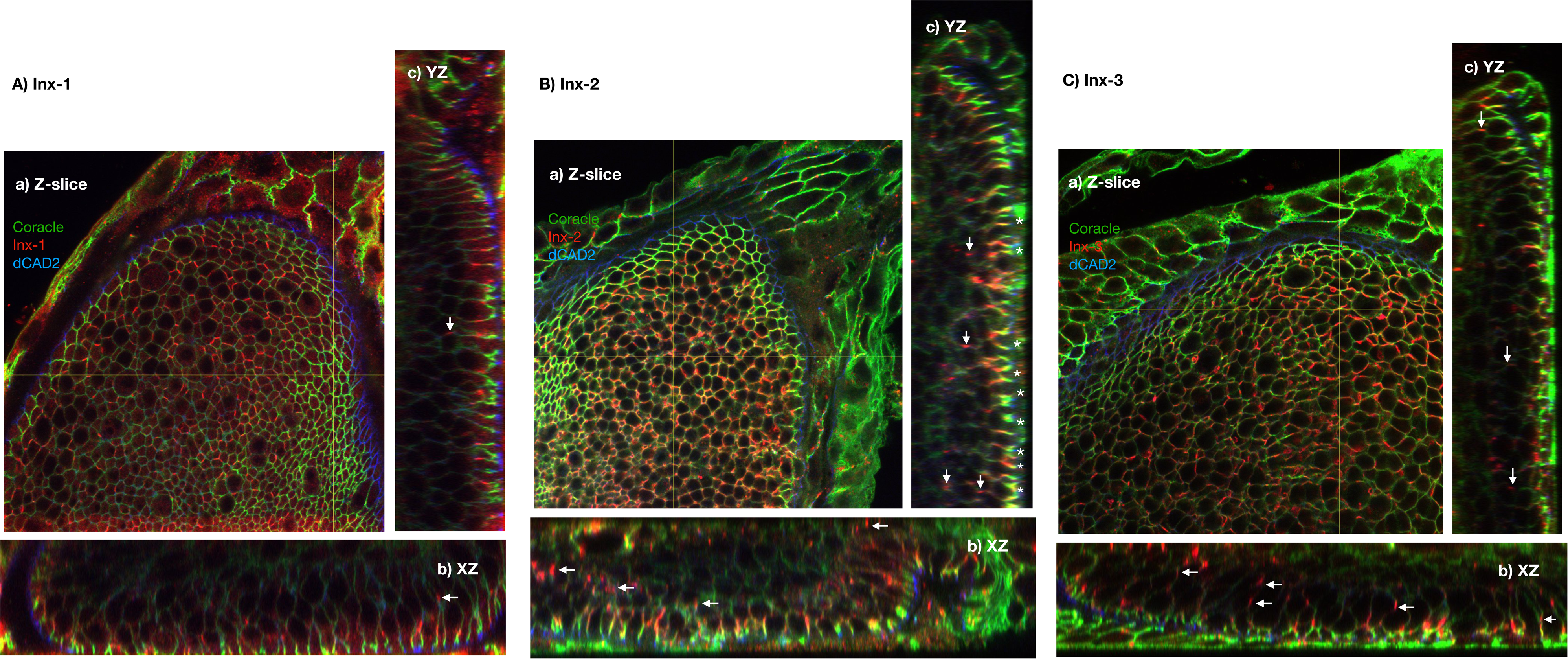
Spatial distribution of innexin-1,-2, and-3 relative to apical junctional markers in disc proper cells. (A–C) Confocal Z-sections **(a, top view)** and orthogonal views **(b, c)** of disc proper (DP) cells show innexin localization (red) in relation to D-Cadherin (blue) and Coracle (green). **(A)** Innexin-1 is positioned sub-apically, beneath adherens and septate junctions, differing from its previously reported basolateral localization in embryonic hindgut and salivary gland epithelia (Bauer *et al*., 2003). **(B)** Innexin-2 displays a similar sub-apical distribution, with partial colocalization with Coracle (**yellow, asterisk**), in contrast to its embryonic localization (Bauer *et al*., 2004). **(C)** Innexin-3 also localizes sub-apically in the notum, though its distribution diverges from earlier mRNA *in-situ* data (Stebbings *et al.,* 2002). All three innexins additionally extend to mid-and baso-lateral domains along the apico-basal axis of disc proper cells (**white arrows in A, B and C**). All panels show magnified views of the notum region.

To further substantiate our findings, we performed tissue-specific RNAi-mediated knockdown of *innexin* genes in the notum region of wing discs using the *pannier-Gal4* (*pnr-*Gal4) driver. The *pnr* gene encodes a GATA family transcription factor that specifies the mediodorsal regions of the thoracic and abdominal segments across embryonic, larval, and adult stages of a fly. During larval development, *pnr* expression in imaginal discs demarcates the presumptive notum region (Calleja *et al*., 2000). To validate the specificity and spatial expression of the *pnr-Gal4* driver prior to knockdown experiments, we generated a *pnr-Gal4, UAS-mCD8::GFP* recombinant fly line. Immunohistochemical staining confirmed GFP expression in the dorsal-most region of the wing disc, consistent with the known activity of the *pnr-Gal4* driver (Figure 5). This validated line was then used to achieve the targeted silencing of *innexin-2* and *innexin-3*.

**Figure 5.**
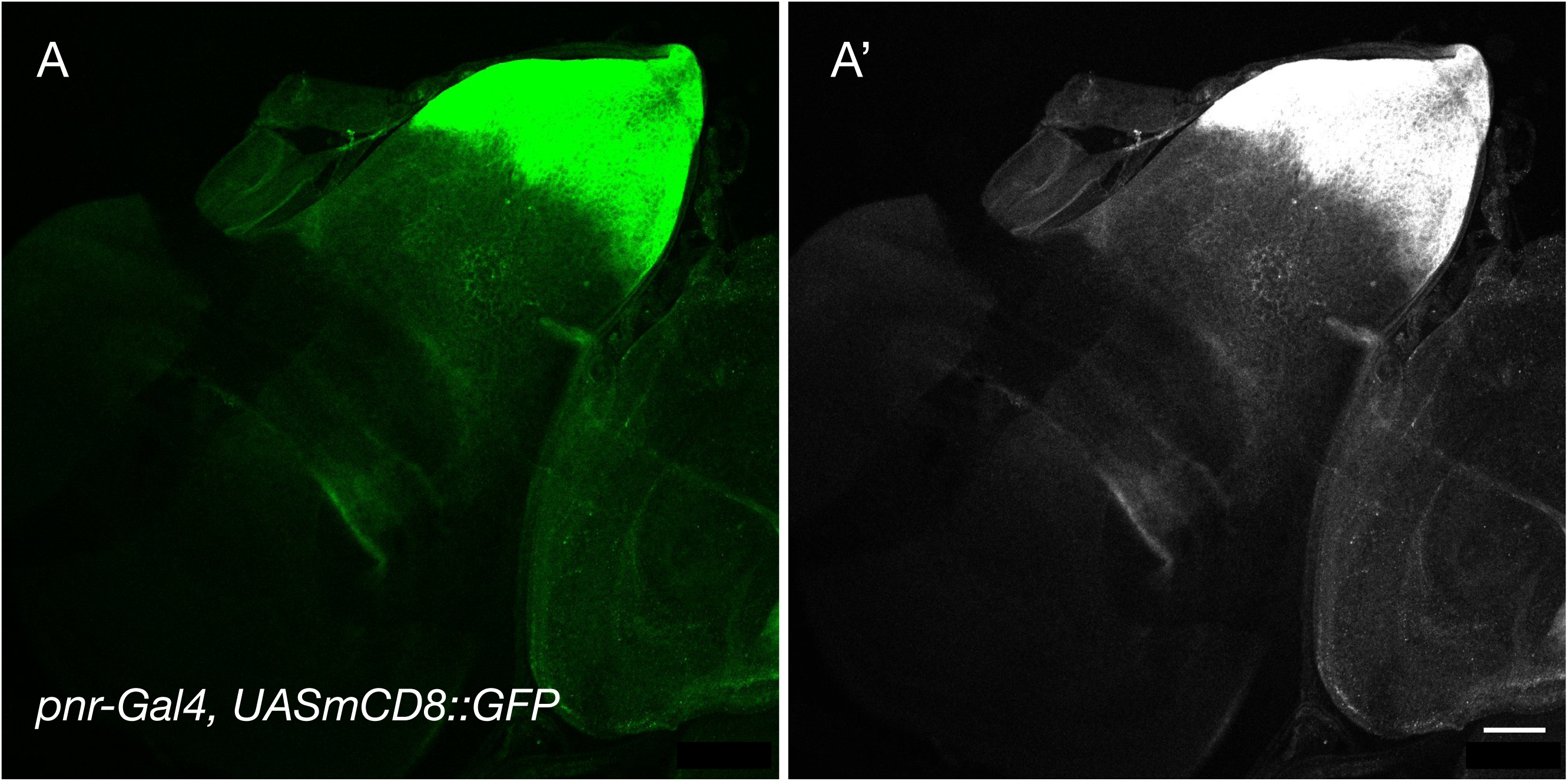
Expression pattern of pannier in wing imaginal discs. Using the recombinant line *pnr-Gal4, UAS-mCD8-GFP/TM6B, Tb*, the pannier expression domain was visualized via GFP, which localized to the dorsal-most region (notum) of the wing imaginal disc. Panels A and A’ show a third instar wing disc stained with anti-GFP antibodies, displayed as a color-composite image (A) and the corresponding grayscale channel (A’). **Scale bars: 50 μm**

For knockdown, we used a *pnr-Gal4/TM6B, Dfd-eYFP, Sb* driver line in combination with either a UAS-RNAi-inx2 or UAS-RNAi-inx3 lines. Thereupon the non-GFP wandering third instar larvae were selected for dissection and immunostaining of wing discs. Genetic silencing of *innexin-2* led to a complete loss of its immunoreactivity within the notum, affecting both disc proper and peripodial epithelial cells. Notably, this knockdown was also associated with a significant reduction in the levels of coracle, a septate junction-associated protein (Figures 6 and 7). These findings are consistent with our observation of innexin-2 colocalization with coracle and suggest that innexin-2 may play a critical role in stabilizing the apicolateral complex comprising adherens, septate, and gap junction components.

**Figure 6.**
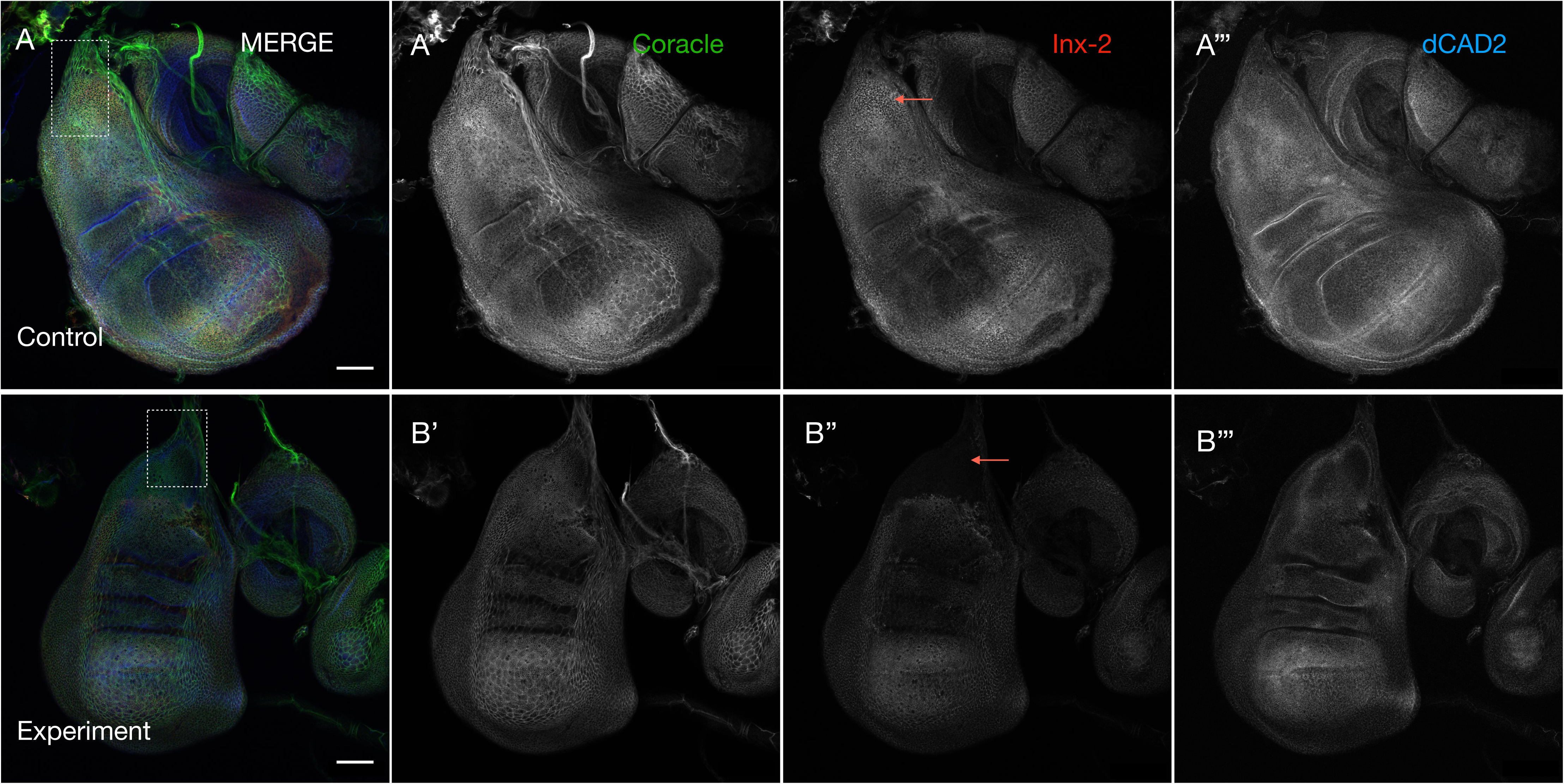
Tissue-specific knockdown of *innexin-2* in the wing discs. RNAi-mediated depletion of *innexin-2* (red) using the *pnr-Gal4* driver results in a pronounced loss of its expression in the targeted region of the wing disc. This knockdown is associated with reduced Coracle (green) levels and noticeable alterations in disc shape and size. **Panels A-A’’’ and B-B’’’** show the control *(pnr-Gal4/+)* and experimental *(pnr-Gal4/UAS-inx2RNAi*) discs. The corresponding grayscale channels for each marker are shown in A’, A’’, A’’’ and B’, B’’, B’’’, respectively. White boxes in panels A and B indicate the *pnr-Gal4* expression domain targeted for knockdown. Red arrows mark the region exhibiting loss of *innexin-2* upon knockdown. **Scale bars: 50 μm**

**Figure 7.**
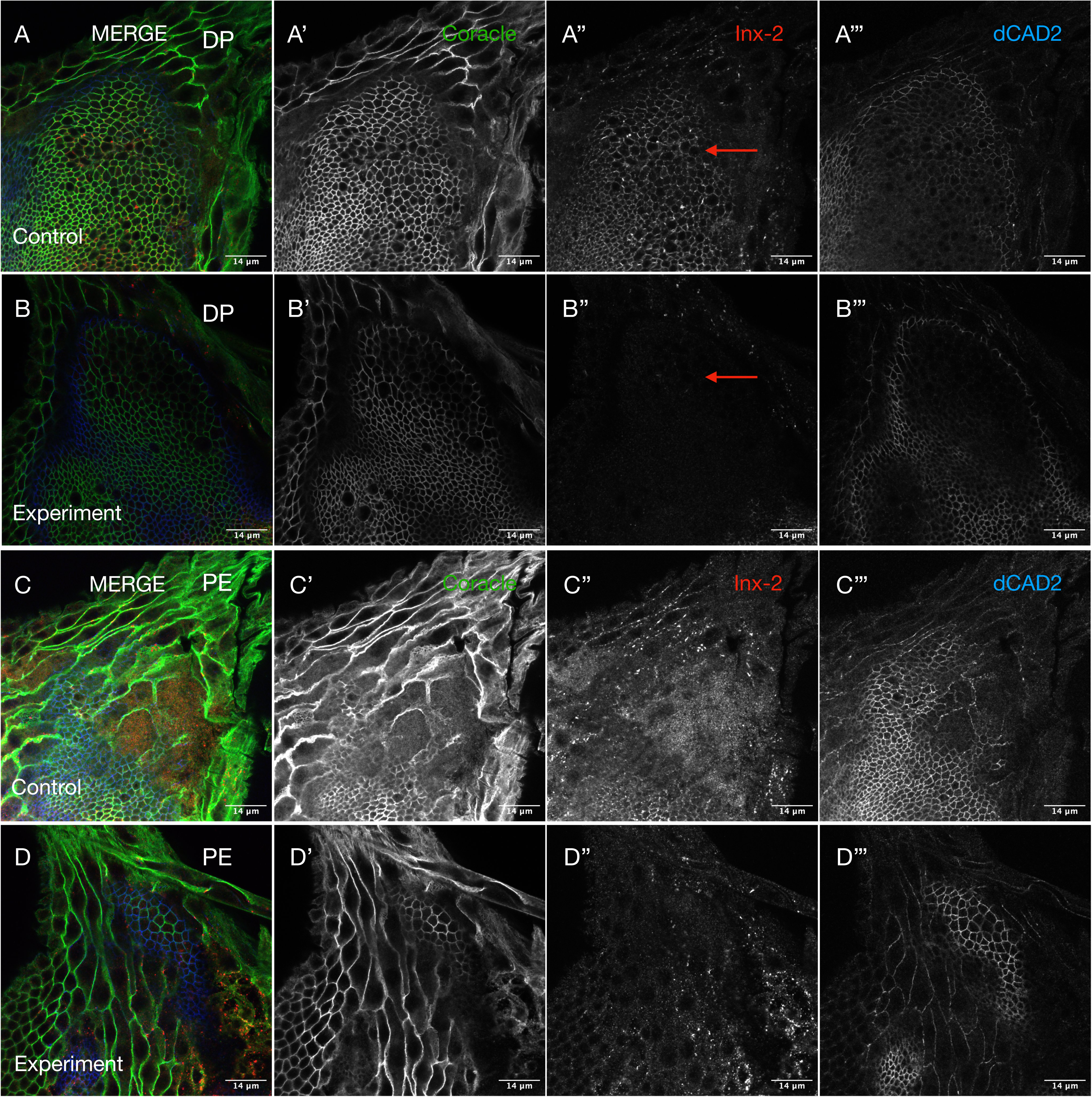
Tissue-specific knockdown of *innexin-2* in the notum region of the wing disc affects expression in both disc proper and peripodial epithelial cells. Panels A–A’’’ and B–B’’’ show the notal region of the disc proper in control (*pnr-Gal4/+*) and knockdown (*pnr-Gal4/UAS-inx2RNAi*) discs, respectively. RNAi-mediated knockdown of *innexin-2* (red) using the *pnr*-Gal4 driver leads to loss of its expression in disc proper cells (compare A’’ and B’’), accompanied by reduced Coracle levels (green; compare A’ and B’). **Panels C-C’’’ and D-D’’’** show the situation with peripodial cells of the control and experimental discs stained with the same markers. Similar to disc proper, loss of innexin-2 expression (C’’ and D’’) and decreased Coracle levels (C’ and D’) are observed. **DP**-Disc Proper and **PE**-Peripodial Epithelia. **Scale bars: 14 μm. Red arrows show the affected region.**

Interestingly, knockdown of *innexin-3* led to a significant reduction in its protein levels within the notum, along with a notable decrease in overall disc size (Figure 8). Both the disc proper and peripodial cells were affected, and the cells appeared more spherical, indicative of cell polarity defects (Figure 9). In addition to these cellular defects, the overall morphology of the wing discs was significantly altered in knockdown specimens, with structural abnormalities extending to the basal regions of the tissue (Figure 10). This phenotype is consistent with our expression data, which show localization of innexin-3 near the basement membrane—a structure essential for tissue organization, connectivity, and mechanical support (Jayadev and Sherwood, 2017).

**Figure 8.**
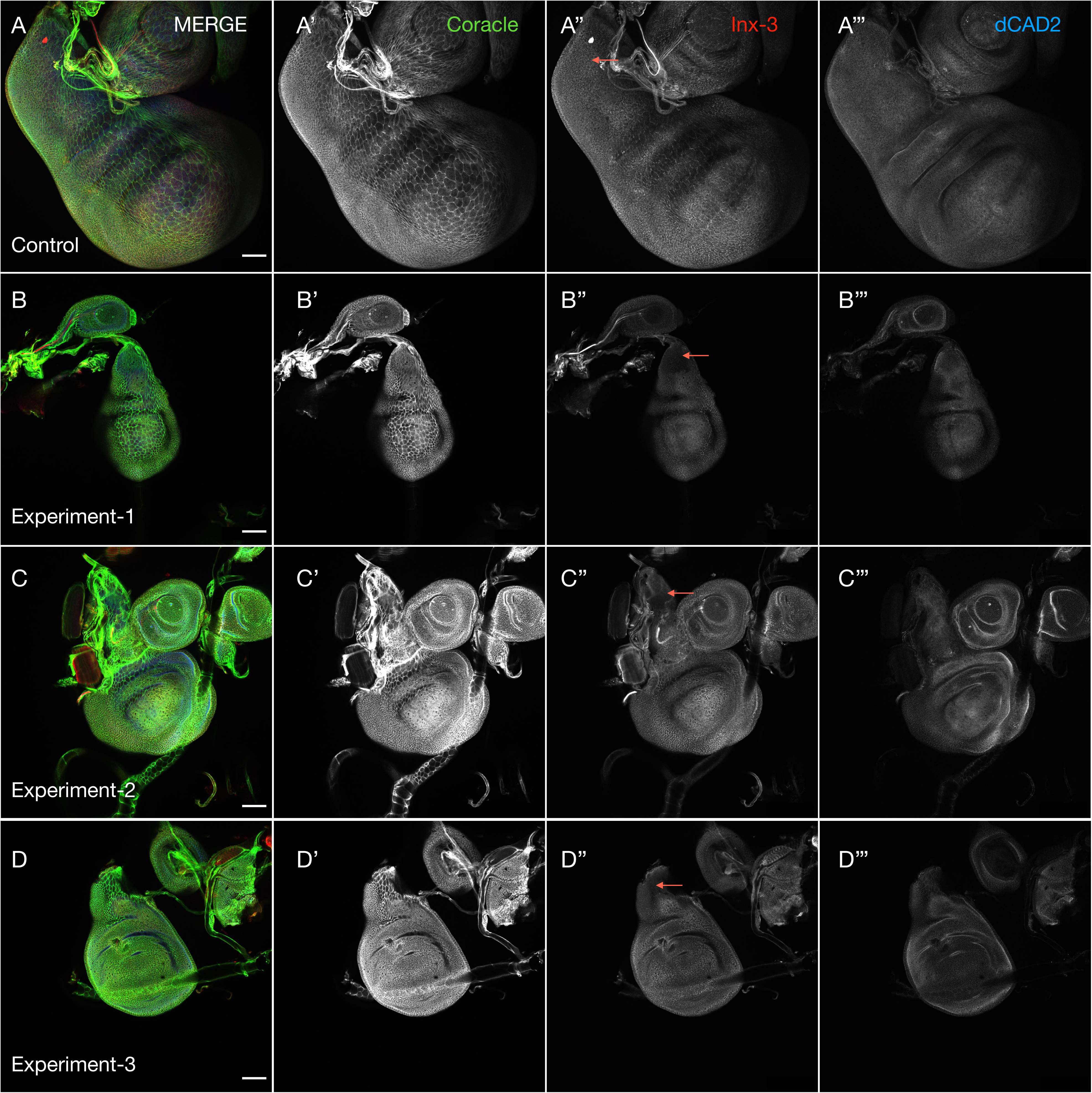
Tissue-specific knockdown of *innexin-3* in the wing discs. RNAi-mediated knockdown of *innexin-3* achieved using the *pannier-Gal4 (pnr-Gal4)* driver resulted in the loss of its expression in the notum region and was accompanied by a reduction in overall disc size. **(A–A’’’)** Control wing disc (genotype: *pnr-Gal4/+*) stained for Coracle (green), innexin-3 (red), and Cadherin (blue). Grayscale images of the individual channels are shown in A’–A’’’ for clarity. **(Panels B, C and D)** Experimental discs (genotype: *pnr-Gal4/UAS-inx3RNAi*) stained with the same markers. Corresponding grayscale images are shown in B’–B’’’, C’– C’’’ and D’–D’’’. A clear loss of innexin-3 expression is observed in the notum region of the experimental discs, as indicated by red arrows. **Scale bars: 50 μm**

**Figure 9.**
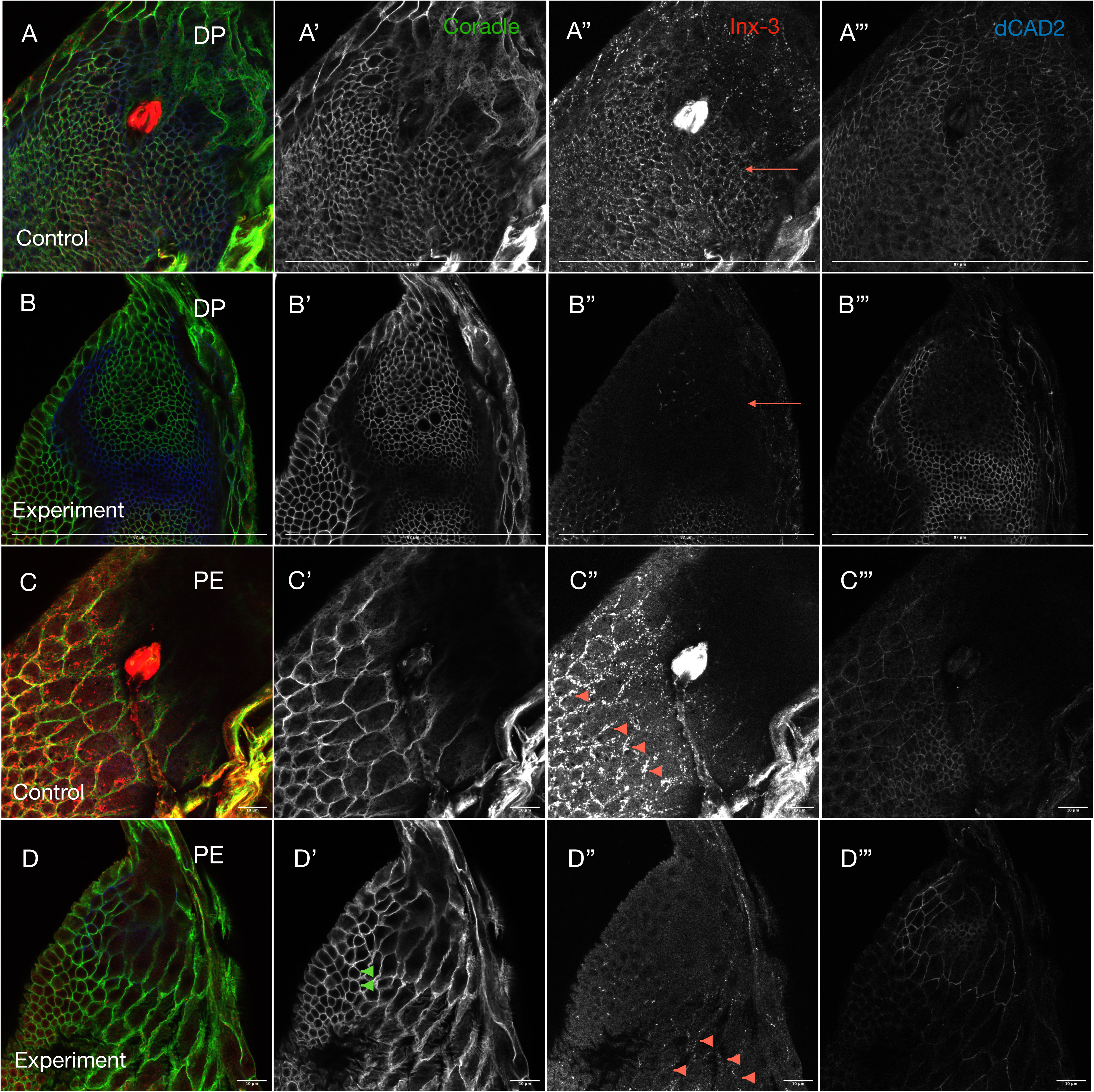
Tissue-specific knockdown of *innexin-3* in the notum region of the wing disc results in its reduced expression in both disc proper and peripodial cells. (A, B) Confocal images of the notum region showing disc proper cells in control **(A)** and experimental **(B)** discs stained for Coracle (green), Innexin-3 (red), and Cadherin (blue). **(A’–A’’’ and B’–B’’’)** Grayscale images of individual channels are shown in **A’–A’’’** and **B’–B’’’** for clarity. Knockdown of *innexin-3* in the experimental discs leads to a marked loss of its expression. **(C, D)** Confocal images of the peripodial cells in the notum region of control (C) and experimental (D) discs stained for the same markers. **(C’–C’’’ and D’–D’’’)** Grayscale representations of individual channels are shown in C’–C’’’ and D’–D’’’. A noticeable reduction in innexin-3 levels at the membranes of the peripodial cells is observed in the experimental discs, compared to controls (the red arrowheads in C’’ and D’’). The peripodial cells in experimental discs also exhibited rounding (green arrowheads in D’), indicative of loss of cell polarity. **DP**-Disc Proper and **PE**-Peripodial Epithelia.

**Figure 10.**
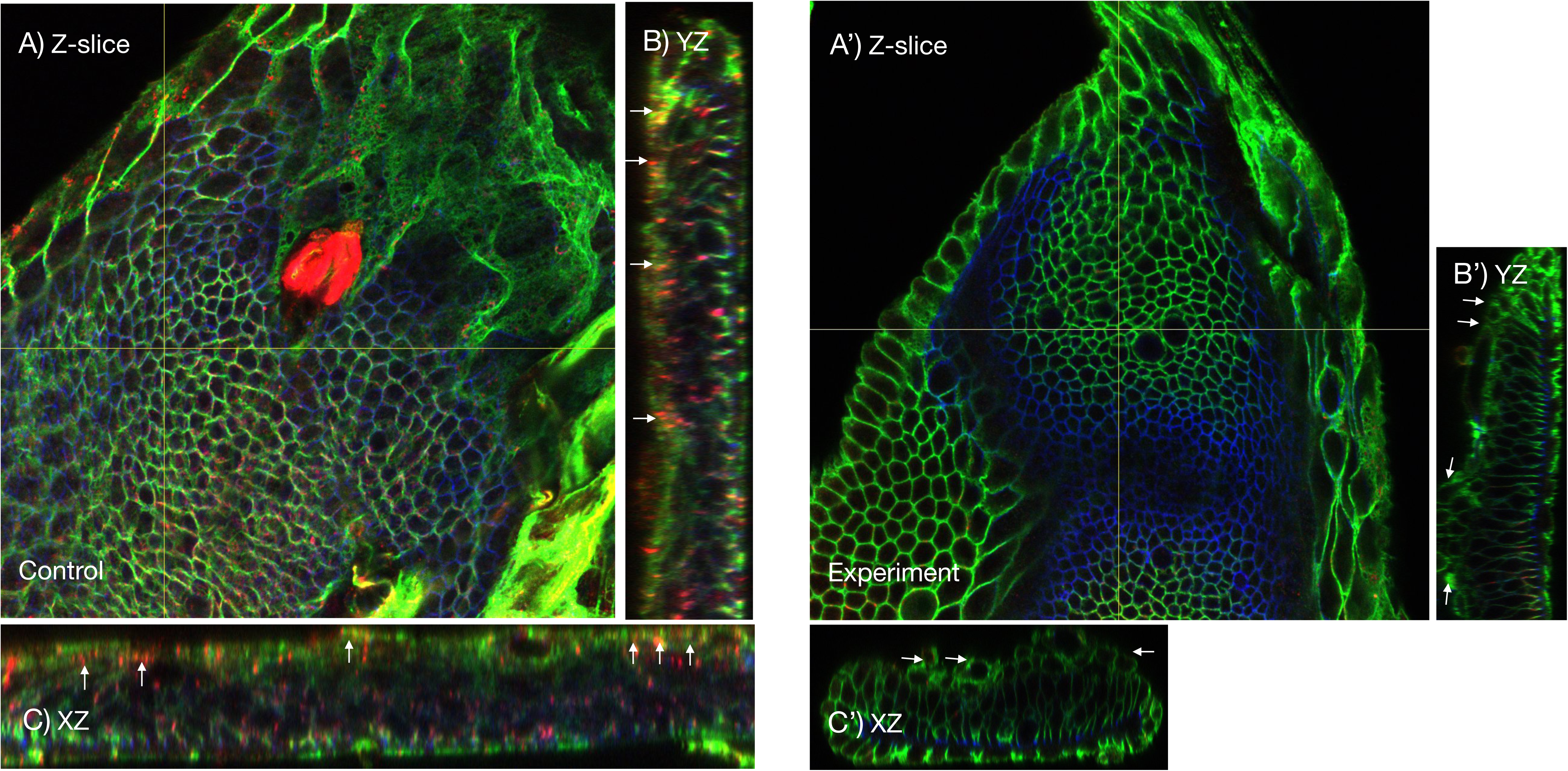
Genetic knockdown of *innexin-3* compromises epithelial membrane integrity towards the basal side of the wing discs. (A, B, and. **C)** display the top (A) and orthogonal (B and C) views of control wing discs (*pnr-Gal4/+*), with white arrows indicating the presence of innexin-3 near the basal side. **(A’, B’, and C’)** show corresponding views of experimental discs (*pnr-Gal4/UAS-Inx3RNAi*), where white arrows highlight the absence of innexin-3 and visible disruptions in the epithelial membrane on the basal side. Additionally, changes in the size and shape of the notum are observed.

The above findings differ from those reported in embryos, where *innexin-2* and *innexin-3* form heteromeric complexes, and *innexin-3* knockdown in the embryonic epithelia phenocopies the *innexin-2 kropf* mutant phenotype (Lehmann *et al*., 2006), suggesting that innexin organization and functional specialization are context-dependent. Furthermore, our study presents a comprehensive visualization of innexin expression within the notum region of the wing disc, confirming their presence in this morphogenetically dynamic domain and proposing a spatial arrangement model based on our current confocal analysis (Figure 11).

**Figure 11.**
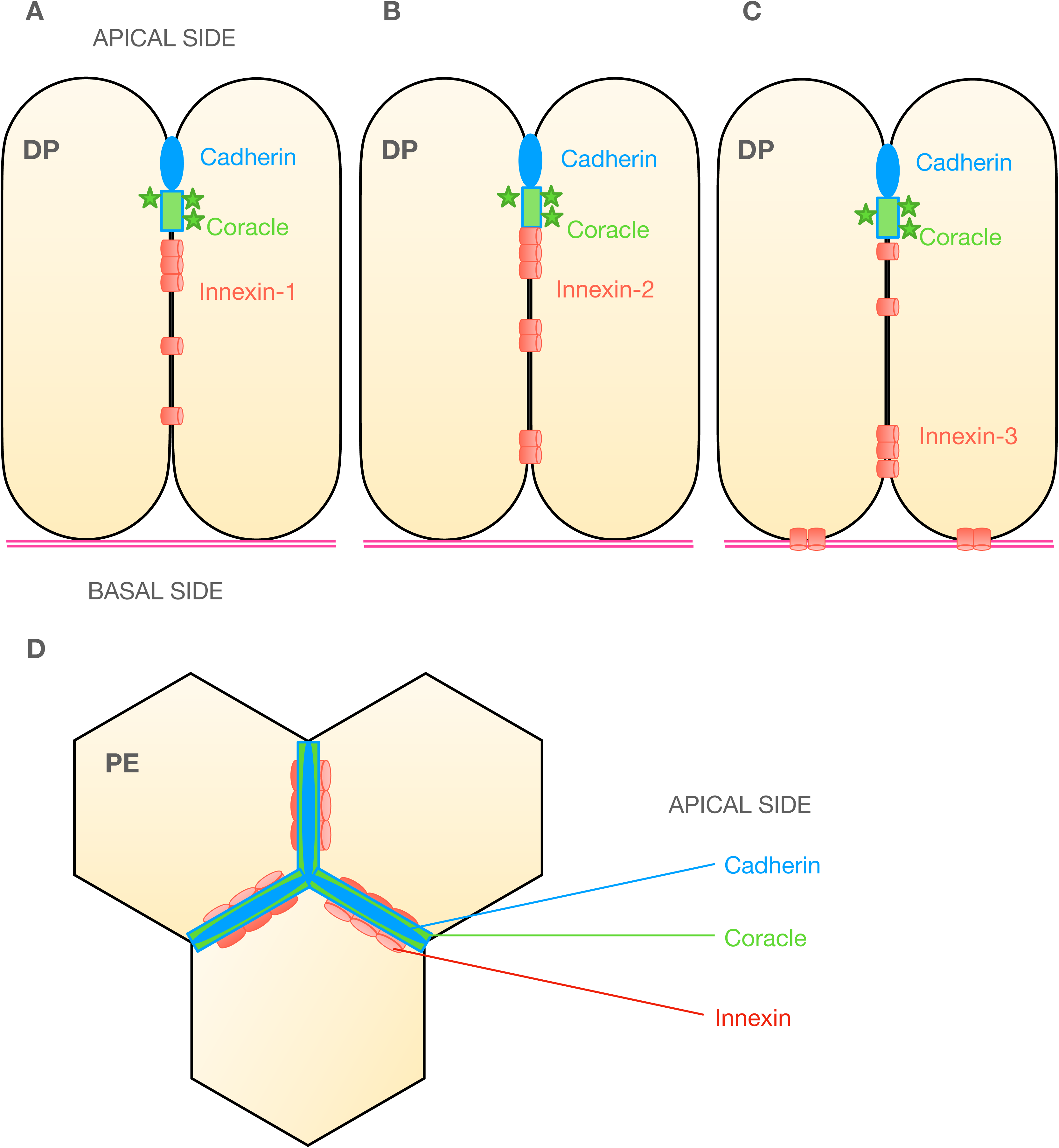
Schematic showing the organisation of gap junction proteins (innexin-1,-2,-3) in disc proper (DP) and peripodial epithelial (PE) cells of the wing disc. (A–C) In DP cells, all innexins localise sub-apically beneath adherens (cadherins) and septate junctions (Coracle). Innexin-2 partially colocalises with Coracle and extends to mid-lateral regions, while innexin-3 also accumulates basolaterally. **(D)** A planar view of the PE cells showing the relative arrangement of cadherin, coracle and innexins from the apical to basal side. While other junctional components extend to tricellular junctions (black arrow), innexins remain largely confined to bicellular interfaces in peripodial epithelial cells.

## Discussion

We specifically report the cellular distribution and arrangement of innexin-1,-2 and-3 in the cells of the wing disc epithelia. Although previous studies have demonstrated the presence and significance of gap junctions in multiple aspects of *Drosophila* wing development, a comprehensive characterization of the cellular distribution and organization of innexins in third instar wing discs is required to offer a definitive spatial context. Our study resolves a key knowledge deficit by providing the protein expression pattern and mapping the cellular localisation and organisation of gap junctions, specifically innexin-1,-2 and-3 in third instar wing discs, providing critical insights that advance current understanding. Our immunohistochemical analyses reveal that all three innexins are broadly expressed across the wing disc, encompassing both epithelial layers, and show continuous as well as punctate membrane localization. This widespread distribution of gap junctions across the wing discs points to their potential involvement in orchestrating morphogenetic events. Specifically, they may facilitate the regulation of epithelial cell shape transitions, such as the flattening of cuboidal cells into squamous cells and the elongation of cuboidal cells into columnar forms, processes that are essential for proper tissue remodelling and wing disc development (McClure and Schubiger, 2005). Moreover, given that the two epithelial layers of the wing imaginal disc, the peripodial epithelium and the disc proper, are known to communicate with each other, and that the peripodial cells respond to cues from the disc proper and vice versa (Pallavi and Shashidhara, 2003; McClure and Schubiger, 2005; Gibson and Schubiger, 2000), to influence organ size, the widespread presence of gap junctions across both layers suggests a potential role for these structures in coordinating growth and regulating tissue size.

Beyond their expression pattern and broad distribution, however, the precise subcellular arrangement of innexins within the wing disc epithelia provide further insight, particularly when compared to their organization in other polarized epithelial tissues such as embryonic epithelium, salivary glands, gut and eye discs. Importantly, we show how the subcellular arrangement of innexins in wing disc cells differ from their organization in these tissues. In salivary glands, for example, gap junctions (innexin-1 and innexin-2) are localized basolaterally and overlap with coracle, a septate junction–associated protein and homolog of vertebrate Protein 4.1; in the hindgut, they are positioned laterally; whereas in embryonic epithelia, innexin-2 localizes primarily apically and colocalizes with adherens junction proteins (Bauer *et al*., 2004; Bauer *et al*., 2005). In contrast, within the disc proper cells of the wing imaginal disc, the membrane proteins are arranged along the apico-basal axis in the order: adherens junction components (cadherins), followed by the septate junction associated protein coracle, and innexins (1, 2, and 3), positioned sub-apically beneath them. All three innexins are primarily localized towards the apical region, with innexin-2 exhibiting partial colocalization with coracle. In addition, innexin-2 and innexin-3 junctions are also detected mid and baso-laterally, with innexin-3 further enriched near the basal side of the disc epithelium. Hence, in contrast to innexin-1, innexin-2 and innexin-3 displayed a more dynamic or broader distribution within the plasma membrane. The distinct distribution of innexin-3 suggests a potential role for innexin-3 in supporting or preserving epithelial cell integrity near the basement membrane, possibly through gap junction mediated interactions with basement membrane components. Furthermore, the heterogeneous localization patterns of innexin-2 and innexin-3 indicate their dynamic behavior within the plasma membrane, in line with live-cell imaging studies in vertebrate systems where connexin hemichannels laterally migrate before assembling into gap junctions with neighboring cells (Lauf *et al.,* 2002). Interestingly, this broad distribution is not observed in eye discs where all three innexins are localised exclusively to the apico-lateral domain. Together, these observations highlight the diverse localization patterns of innexins across epithelial systems and underscore their context-dependent roles in cellular communication and tissue organization.

In the peripodial epithelium, innexins were detected in distinct punctate patterns along the cell membranes, reminiscent of previously reported punctate localizations of innexins in embryos (Bauer *et al*., 2004), ovaries (Bohrmann & Zimmermann, 2008), and eye discs (Richard *et al*., 2017), as well as connexins in vertebrate tissues (Rash *et al*., 2001; Zhang & Hill, 2005). Such punctate distributions are generally interpreted as clusters of gap junction proteins that assemble into specialized membrane domains mediating intercellular communication. The variation in punctate organization among different innexins likely reflects differences in their molecular properties and functioning. The bright, dense puncta detected with innexin-2 staining may correspond to actively engaged junctional complexes, whereas the enrichment of innexin-1 and innexin-3 at bicellular junctions suggests spatially restricted trafficking or signalling events.

Additional validation of these expression patterns was achieved through tissue-specific RNAi-mediated knockdown of *innexins* in the notum region. In the wing imaginal disc, the notum plays a critical role—it connects the disc to the body wall epithelium and undergoes dynamic movements during metamorphosis to facilitate dorsal thorax closure. It also harbours the adult muscle progenitors (AMPs) and trachea on the basal side, highlighting its functional and structural complexity. Genetic knockdown of *innexin-2* and *-3* within this region produced distinct phenotypes, suggesting that co-expressed innexins in the same region may participate in separate signalling pathways and perform specialised functions. Preliminary analyses revealed that downregulation of *innexin-2* led to a reduction in its expression within the notum, accompanied by decreased levels of coracle and changes in wing disc morphology, underscoring the role of innexin-2 in epithelial morphogenesis and the maintenance of tissue integrity (Bauer *et al.,* 2004). In contrast, silencing of *innexin-3* resulted in its selective loss from the notum along with a decrease in overall disc size. Additionally, *innexin-3* knockdown was associated with structural defects near the basement membrane, which may underlie the observed changes in cell shape and the compaction of the tissue. These findings also highlight how the localised absence of a specific innexin can differentially influence the architecture and function of the entire tissue. The presence of gap junctions in the notum region opens up intriguing possibilities for further research, particularly in understanding how the distinct roles of individual innexins influence cellular function, tissue development, and overall organismal behavior. The observed differences in their localization, whether distributed across distinct or partially overlapping membrane domains, may help explain both the functional redundancies and unique roles attributed to different innexins. In summary, the manuscript reports characterisation of innexin expression in third instar wing imaginal discs, with implications for understanding the functional diversity and spatial dynamics of gap junctions in development.

## Acknowlegements

We thank the members of the VijayRaghavan Lab at NCBS, India, for their valuable support in the initial design and formulation of this project. The author, Spraha Bhandari, sincerely acknowledges Prof. K. VijayRaghavan for hosting her EMBO scientific exchange application and providing access to laboratory facilities, and the NCBS Confocal Data Collection Facility for their support and resources, which were essential for the data collected in this work. She further expresses her gratitude to Prof. Satyajit Mayor for supporting her ICMR application as signing authority in his capacity as Director.

## Author Contributions

S.B - conceptualisation, funding acquisition, fly experiments and data collection, writing and reviewing.

A.C - scientific inputs and generation of recombinants.

F.E - scientific inputs and support in funding acquisition.

R.B - valuable scientific inputs and feedback towards conceptualisation, provision of reagents, help with funding acquisition and grant writing, manuscript writing and reviewing.

## Use of AI technology

During the preparation of the manuscript, ChatGPT was used to enhance readability and comprehension of the work. The authors subsequently reviewed and edited the content as necessary and take full responsibility for the final version of the publication.

